# Non-invasive quantification of hepatic necrosis from circulating alanine aminotransferase kinetics in acetaminophen-treated mice in vivo

**DOI:** 10.64898/2026.09.11.750627

**Authors:** Chris Humphries, Jennifer A. Cartwright, Rhona Aird, Maria Elena Candela, Anuruddika J. Fernando, Philip Starkey Lewis, Janet Man, Charlotte Potter, Kathleen M. Scullion, Justyna Cholewa-Waclaw, James W. Dear, Stuart J. Forbes

## Abstract

Histological necrosis is the reference measure of hepatotoxicity but can only be obtained at terminal cull. A time course requires independent cohorts at every timepoint. Circulating alanine aminotransferase (ALT) can be sampled repeatedly in the same animal, and is released in liver injury. We asked whether serial ALT kinetics can be used to estimate histological necrosis in vivo. Forty-five fasted twelve-week-old male C57BL/6J mice received a single intraperitoneal dose of 350 mg/kg acetaminophen (paracetamol). Plasma ALT and microRNA-122 (miR-122) were measured serially from baseline to 48h as cumulative area-under-the-curve (AUC) to cull, against centrilobular necrosis on haematoxylin and eosin sections as reference (range 0 to 59%, mean 32%). A generalised additive model of necrosis on cumulative ALT AUC and time since dosing predicted necrosis with a leave-one-animal-out cross-validated mean absolute error of 7.7% necrotic area (95% CI 5.6 to 10.0; n=49). A panel of traditional regression and machine-learning models all gave equal or larger error, and adding an additional biomarker or regeneration information did not improve prediction. Serial ALT kinetics therefore provide a calibrated, longitudinal measure of hepatic necrosis in vivo and give a more stable estimate of within-group variance for study planning, while supporting reductions in animal use, because one serially-sampled cohort can replace separate cohorts at each timepoint.

## Introduction

Quantitative histopathology is the reference measure of organ injury in preclinical toxicology and can typically only be obtained at terminal cull. Alanine aminotransferase (ALT) is released into the circulation as hepatocytes die in acute liver injury (Starkey Lewis et al. 2011; McGill et al. 2012; Antoine et al. 2013), but is typically only considered supporting information, as values reflect simultaneous release, ongoing injury and clearance. Kinetic information with potential to resolve this exists in any serially-sampled study and is discarded when single values are reported. We asked whether serial ALT quantification over time since dosing is sufficient to estimate histological necrosis in a previously-described mouse model of acetaminophen (paracetamol, APAP)-induced liver injury (Starkey Lewis et al. 2020; Sherman and Goessling 2024). We assessed the performance of multiple modelling approaches, and the potential impact on sample variability and reduction in animal use.

## Materials and methods

Procedures were approved by the University of Edinburgh Animal Welfare and Ethical Review Body (093-LFR-24) under the Animals (Scientific Procedures) Act 1986 and follow ARRIVE 2.0 (Percie du Sert et al. 2020). After a 14h fast, 45 twelve-week-old male C57BL/6J mice (Charles River, UK) received a single intraperitoneal dose of 350 mg/kg APAP in warm sterile saline; five fasted, undosed control mice were culled at t=0 (0% necrosis). This dose was selected because it produces reproducible submaximal centrilobular necrosis within a UK moderate-severity limit (Starkey Lewis et al. 2020). One mouse found dead at 7h was excluded and seven reaching a humane endpoint off schedule were retained at the nearest timepoint, giving n=49 across seven timepoints to 48h.

Serial 40 µL tail-vein samples were taken at baseline and at 8, 16, 24, 36 and 48h until cull. ALT and miR-122 were measured blind to group and integrated to cumulative area-under-the-curve (AUC) by the trapezoidal rule; necrosis was quantified blind from whole-slide haematoxylin and eosin images by trainable tissue segmentation, as described previously (Humphries et al. 2026). Necrosis was modelled on cumulative ALT AUC and time since dosing by a generalised additive model (GAM), one penalised smooth per predictor on the square-root response scale (Wood 2017), evaluated by leave-one-animal-out cross-validation with smoothing parameters selected inside each training fold and summarised as mean absolute error (MAE) with a 10,000-sample bootstrap 95% confidence interval. Every comparator in Supplementary Table S1 was evaluated identically. The Supplementary Material gives the full husbandry, allocation, welfare and assay-performance detail (Section 1); biomarker AUC construction, model specification and response-scale checks (Section 2); full comparator panel with confidence intervals (Section 3); regeneration analyses (Section 4); study-design, variance and animal-number derivations (Section 5); and per-animal data for all 44 dosed animals (Section 6).

## Results and discussion

Necrosis spanned 0% in the fasted controls to 59% (mean 32%). Single ALT readings reflected necrosis poorly: two dosed animals with near-identical ALT to 16h reached 55% and 18% necrosis (Supplementary Fig. S1).

However, the GAM on cumulative ALT AUC and time (Fig. 1a) predicted necrosis with a cross-validated MAE of 7.7% necrotic area (95% CI 5.6 to 10.0; n=49; Fig. 1c); predictive R² was 0.49 and Bland-Altman bias 0.7% (Bland and Altman 1986; Supplementary Fig. S2), supporting use of the model for group-level inference rather than as an individual-animal assay. The GAM response surface demonstrated expected kinetics: at a given cumulative exposure, predicted necrosis was highest early and declined with time, because later AUC increasingly reflects retained, slowly-cleared enzyme rather than fresh cell death.

**Fig. 1.**
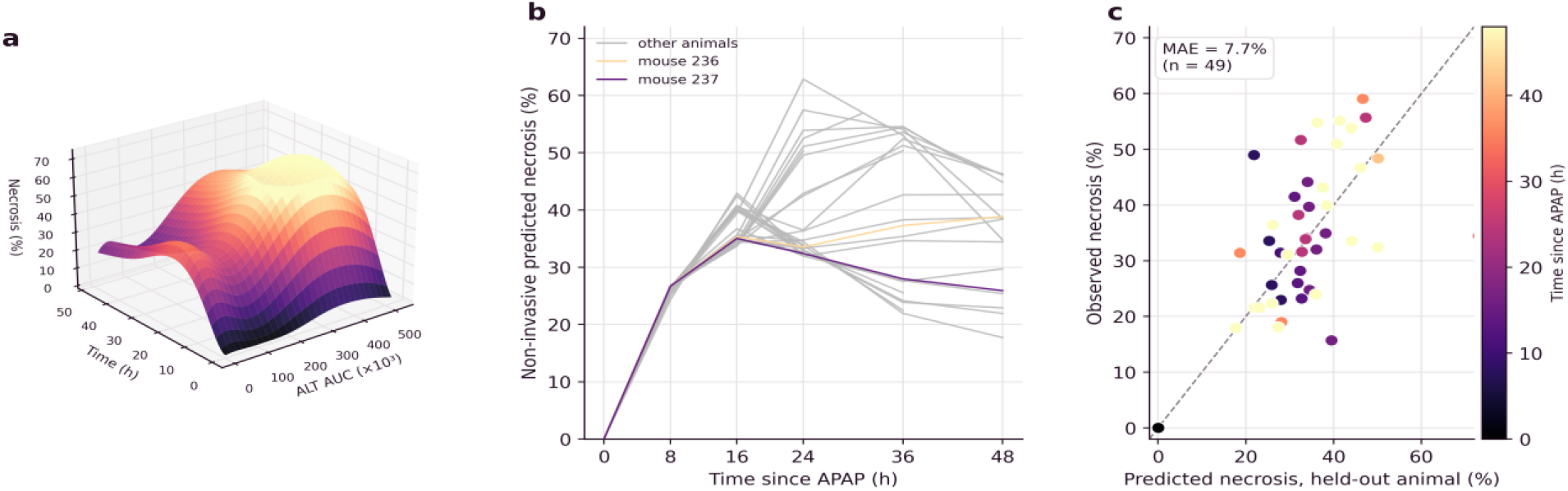
Virtual histology from serial ALT kinetics in acetaminophen-treated mice. **a** Generalised additive model surface: predicted centrilobular necrosis over cumulative plasma ALT AUC and time since dosing. The surface is interpretable only within the range of the observed data. **b** Longitudinal read-out: the model applied along each animal’s serial ALT AUC trajectory returns a necrosis estimate at every sampling point in the living animal. The two example animals with near-identical early ALT (Supplementary Fig. S1) are highlighted. **c** Leave-one-animal-out cross-validated prediction against observed necrosis (mean absolute error 7.7%, n=49); the dashed line is identity and points are coloured by time since dosing.

Because model inputs are obtained serially, necrosis can be estimated at each sampling point in living animals rather than only at cull (Fig. 1b). This signal can be identified in every modelling approach we studied: from log-log regression to a neural network, confidence intervals overlapped the GAM (Supplementary Table S1; Figs. S3, S4). The GAM was retained for interpretability and the ability to make predictions outside of observed data. Including additional biomarkers did not improve performance: miR-122 was worse in isolation, collinear with ALT (r=0.82)and combining both biomarkers did not help (Supplementary Fig. S5). Incorporating regenerative signals also did not improve prediction, nor did incorporating the time of peak ALT (Supplementary Section 4; Fig. S6).

Serial sampling stabilised the sample variance which studies are powered on: the within-timepoint coefficient of variation of cumulative ALT AUC remained stable (34 to 38%) in the serially-bled cohort but was unstable (15 to 45%) across the independent cull groups (Supplementary Section 5; Fig. S7). Additionally, because one serially-sampled cohort replaces one cohort per timepoint, a time course at matched precision needs 50% to 80% fewer animals for two to five timepoints. This would translate to 40 to 48 instead of 144 animals for a two-group, three-timepoint study powered on a 10% necrosis reduction.

We note that the calibration assumes injury releases ALT in proportion to hepatocyte death, so predominantly apoptotic or biliary injury would be underestimated. Our model was derived in one mechanism, dose, sex and strain, and transfer to other hepatotoxicants, doses and to females requires testing. The analyses supporting each of the points above, including the model-specification checks, the full comparator panel, the regeneration data and the sample-size derivations, are set out in the Supplementary Information. However, the principle translates to any tissue injury that releases a tissue-selective biomarker in proportion to the damage. External validation across injury mechanisms, and prospective use with a retained terminal-histology subset, are the necessary next steps.

## Supporting information

ARRIVE

## Statements and Declarations

### Ethics approval

All animal procedures were approved by the University of Edinburgh Animal Welfare and Ethical Review Body (project licence 093-LFR-24) and conducted under the Animals (Scientific Procedures) Act 1986. No human participants were involved, so consent to participate and consent to publish are not applicable.

### Author contributions

Conceptualization, software, data curation, visualization and project administration: CH. Methodology, formal analysis and writing of the original draft: CH, JAC. Investigation: CH, JAC, RA, MEC, AJF, JM, CP, KMS, PSL, JCW. Resources and supervision: SJF. Review and editing: all authors. Funding acquisition: JWD, SJF.

### Data availability

Per-animal data are provided in the Supplementary Information. The analysis and verification code are available from the corresponding author on reasonable request.

## Acknowledgements

We thank the Bioresearch and Veterinary Services staff at the University of Edinburgh.

