## supplemental for "Non-invasive quantification of hepatic necrosis from circulating alanine aminotransferase kinetics in acetaminophen-treated mice in vivo"

### Electronic Supplementary Material

This document provides the supporting methods, model specification and verification, the comparator panel, the regeneration analyses, the animal-number derivation, seven supplementary figures and the per-animal data table. Section numbers correspond to the ESM cross-references in the main text.

#### Section 1. Animals, study design and endpoint measurement

##### *1.1 Allocation, group sizes and welfare*

Fifty twelve-week-old male C57BL/6J mice (Charles River, UK) were housed in groups of five on a 12 h light/dark cycle with free access to water. After a 14 h fast, 45 mice received a single intraperitoneal dose of 350 mg/kg acetaminophen (APAP) in warm sterile saline, following the protocol of Starkey Lewis et al. (2020). Five fasted-only mice were culled at t = 0 without dosing to confirm baseline ALT validity (paired change from baseline p = 0.60; 0% necrosis).

Allocation to cull timepoint was sequential rather than randomised, because mice were co-housed by timepoint to maintain social housing and reduce isolation stress. ALT and necrosis measurements were blinded to group. One dosed mouse found dead at 7 h was excluded. Seven mice reached a humane endpoint, assessed against protocol-specific welfare scoring criteria (Cartwright et al. 2025), outside scheduled times and were retained at the nearest scheduled group (three culled at 11, 13 and 14 h retained at 13 h; two culled at 31 and 42 h retained at 36 h). Final group sizes were: 0 h control n = 5; 8 h n = 5; 13 h n = 5; 16 h n = 6; 24 h n = 6; 36 h n = 5; 48 h n = 17. The 48 h group was enlarged to enable a planned comparison of serial sampling against serial cull on group variability. Forty-four dosed mice entered the models, giving n = 49 analysed.

##### *1.2 Assay performance*

Plasma ALT was measured by a blinded laboratory service using the Bergmeyer method (intra-assay coefficient of variation < 4%; inter-assay < 8%). miR-122 was quantified by qPCR normalised to a miR-39 pre-extraction spike-in. Serial 40 µL tail-vein microsamples were taken at baseline and at 8, 16, 24, 36 and 48 h until cull, with a terminal cardiac sample at cull.

##### *1.3 Necrosis quantification*

At cull, livers were formalin-fixed, paraffin-embedded, sectioned at 4 µm and stained with haematoxylin and eosin. Percentage centrilobular necrosis was quantified from whole-slide images acquired at 20x magnification across at least ten regions of interest per animal by a blinded study team member using InForm trainable tissue-segmentation software (Akoya Biosciences), by the segmentation approach described in Humphries et al. (2026).

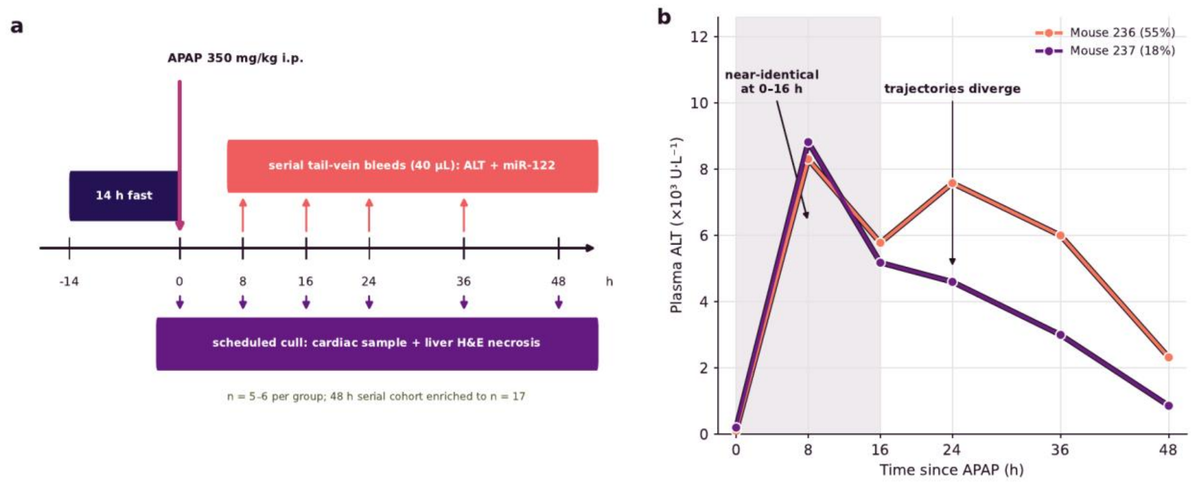


Fig. S1 Study design and rationale. a Study timeline: mice were fasted for 14 h, dosed with 350 mg/kg APAP intraperitoneally, then bled serially for ALT and miR-122 while groups were culled at scheduled times for reference haematoxylin and eosin necrosis. b Why a single timepoint misleads: two animals (mice 236 and 237) with near-identical plasma ALT from 0 to 16 h follow divergent later trajectories and reach necrosis of 55% and 18%, so cumulative kinetics rather than any single reading carry the information. The later divergence likely reflects secondary inflammation driving a rise in ALT for mouse 236 after 16 h.

#### Section 2. Biomarker AUC and model specification

##### *2.1 Biomarker AUC*

Cumulative biomarker exposure to cull was computed by the trapezoidal rule across the serial timepoints available for each animal. For miR-122, qPCR cycle-threshold values were first linearised as 2 to the power of minus ΔCt (ΔCt normalised to the spike-in reference), because the inverse-logarithmic Ct scale is not additive and an AUC computed on raw Ct is not meaningful. Each animal's miR-122 values were mapped onto its true ALT sample-time spine before integration.

##### *2.2 Model specification and cross-validation*

The headline model is a generalised additive model (GAM) with a separate penalised smooth for cumulative ALT AUC and for time since dosing, summed on the square-root response scale, fitted by restricted maximum likelihood (R package mgcv). Predictions were back-transformed by squaring and truncated at zero. Performance was estimated by leave-one-animal-out cross-validation (LOOCV): each animal was predicted from a model refitted on the remaining animals, with the smoothing parameters selected within each training fold, and the mean absolute error (MAE) computed on the natural necrosis scale with a bootstrap 95% confidence interval. A log-log regression comparator, a straight-line fit on logarithmic axes, was used as described in Humphries et al. (2026). The whole analysis was implemented in R (mgcv) and independently re-implemented in Python (pyGAM, scikit-learn); the two agree on all out-of-sample metrics reported here. This is a check on the analysis, not on the experiment, which was performed once.

##### *2.3 Response-scale choice*

The square-root response scale was checked rather than assumed. On the leave-one-animal-out basis, identity, square-root and log response scales gave cross-validated MAE of 7.75%, 7.67% and 8.68% respectively, and the square-root scale did not reduce residual heteroscedasticity relative to the identity scale (correlation of absolute residual with fitted value 0.23 versus 0.21, both non-significant). The square-root scale is retained because it constrains the back-transformed prediction to be non-negative at no measurable cross-validated cost; the identity scale would serve equally well.

##### *2.4 Biological rationale for the time term*

The ALT AUC to necrosis slope decays with time since dosing. Early after injury, synchronous hepatocyte lysis releases ALT in steep proportion to necrosis. Later, accumulated AUC increasingly reflects retained, slowly-cleared enzyme, clearance becomes non-negligible, necrosis becomes more diffuse, and regeneration begins, all of which flatten the apparent relationship. Including time as a continuous covariate lets the model absorb this, and is why a single-timepoint biomarker value is insufficient.

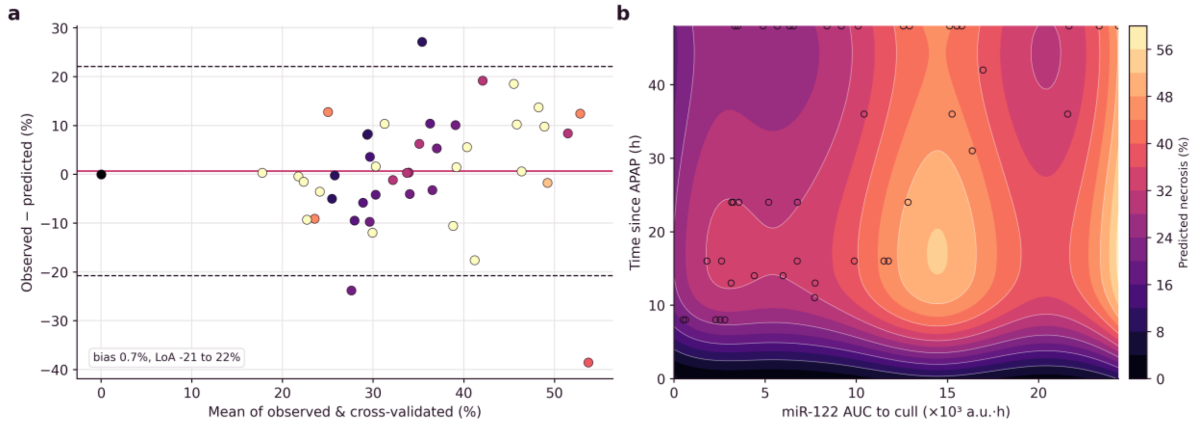


Fig. S2 Model agreement and the miR-122 surface. a Bland-Altman agreement of the ALT model: between observed necrosis and the GAM leave-one-animal-out prediction from ALT (bias 0.7%, limits of agreement minus 21 to 22%). b miR-122 virtual-histology surface: predicted necrosis over cumulative miR-122 AUC and time since dosing.

#### Section 3. Model comparison and specification robustness

All models were compared on identical leave-one-animal-out cross-validation, reporting MAE in percentage necrotic area. GAMs used a square-root response; black-box learners used scikit-learn defaults with standardised inputs. On this basis the additive GAM gave 7.67%, the log-log regression 9.79% and a plain linear fit (necrosis on AUC and time) 10.85%: the flexible model improves out-of-sample error by approximately 21% over the log-log regression and approximately 29% over the plain linear fit. No black-box learner materially improved on the additive GAM (Table S1).

**Table S1** Cross-validated error by model (ALT AUC and time as inputs). Leave-one-animal-out cross-validation with 10,000-sample bootstrap 95% confidence intervals; n = 49 throughout except the log-log regression, which requires log(necrosis) and is fitted on the 44 dosed animals. All intervals overlap the additive GAM, so no comparator is distinguishable from it at this sample size.

| Model | Type | LOOCV MAE (%) | 95% CI |
|---|---:|---:|---:|
| Linear regression | white-box | 10.85 | 8.8 to 13.0 |
| Log-log regression | white-box | 9.79 | 7.7 to 11.9 |
| GAM (additive, headline) | grey-box | 7.67 | 5.6 to 10.0 |
| GAM (tensor interaction) | grey-box | 7.94 | 5.8 to 10.4 |
| Random forest | black-box | 8.07 | 6.3 to 9.9 |
| Gradient boosting | black-box | 8.29 | 6.5 to 10.2 |
| SVM (RBF) | black-box | 8.29 | 6.7 to 10.0 |
| k-NN (k = 5) | black-box | 7.76 | 6.2 to 9.4 |
| Neural net (MLP) | black-box | 12.22 | 9.9 to 14.7 |

k-NN was the closest black-box comparator but, unlike the GAM, cannot provide smooth predictions and degrades where data are thin, so it does not support the continuous surface and serial trajectories the read-out requires.

##### *3.1 GAM specification robustness*

Cross-validated MAE was also stable across the specification of the GAM itself: a tensor-product interaction gave 7.9% at basis dimension (6,5) and 7.5% at the larger mgcv basis (10,9); a quadratic-order tensor gave 8.2%. Only an over-small (4,4) basis underfitted, at 8.6%. The result therefore does not depend on the specific smoother or basis dimension within a sensible range, and the additive form is retained as the most parsimonious specification that achieves the best error.

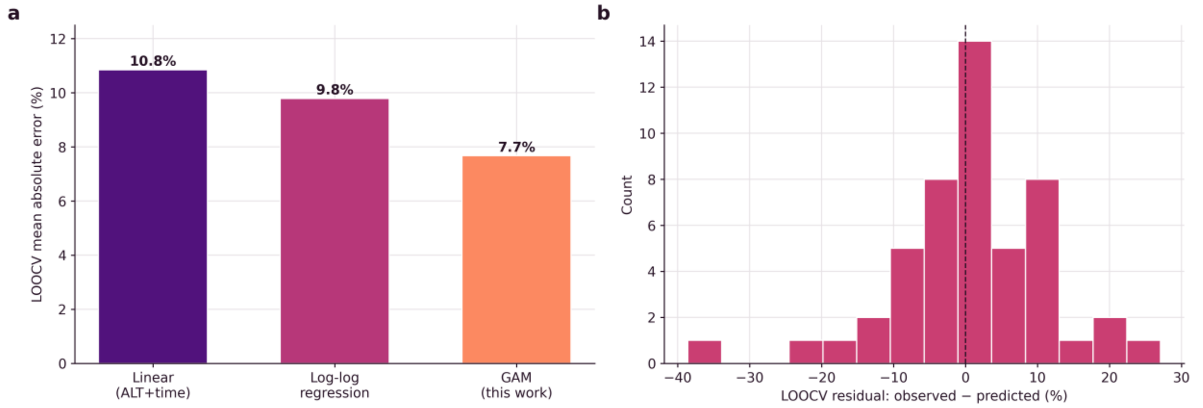


Fig. S3 Model comparison and residuals. a Cross-validated MAE for the plain linear fit, the log-log regression and the GAM. b Distribution of GAM leave-one-animal-out residuals.

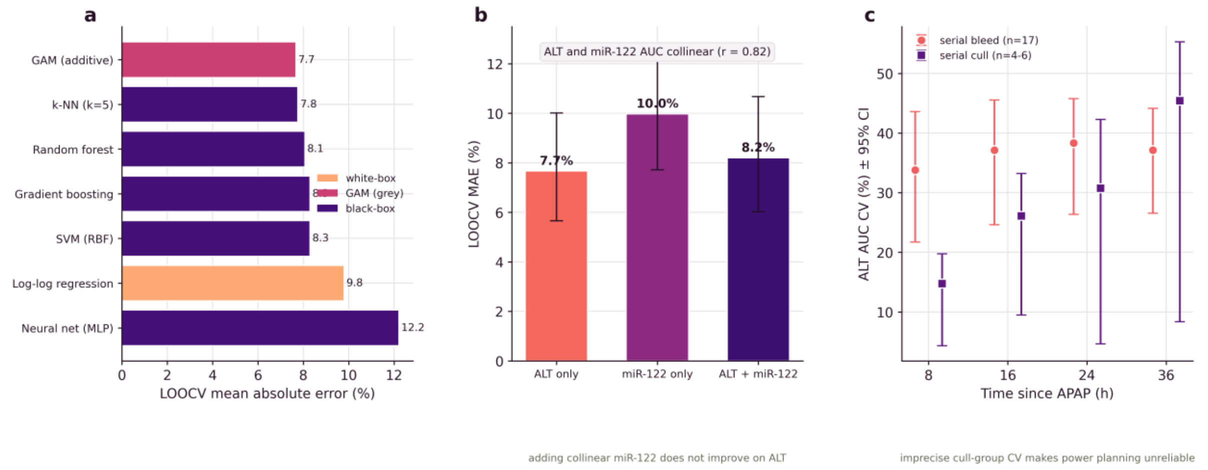


Fig. S4 Model selection and robustness. a Accuracy against interpretability: cross-validated error by model, shaded by interpretability class (white-box methods are directly interpretable, black-box methods are not, and GAM interpretability sits between): the GAM matches or betters the black-box learners. b Combining ALT and miR-122 does not improve on ALT alone; the two AUCs were collinear (Pearson r = 0.82). c Bootstrap 95% intervals for the ALT AUC coefficient-of-variation estimate: the small cull groups give imprecise variance estimates and the serial-bleed design a stable one. Error bars in b and c are 95% bootstrap confidence intervals.

#### Section 4. Biomarker integration and the regeneration axis

Cumulative ALT AUC and cumulative miR-122 AUC were strongly collinear (Pearson r = 0.82), so adding miR-122 to ALT contributes little independent information. Across all n = 49 animals with validated miR-122 AUC, combining the two did not reduce error and slightly increased it under the GAM (Table S2).

**Table S2** Biomarker combination, leave-one-animal-out cross-validation.

| Model inputs | LOOCV MAE (%) | 95% CI |
|---|---:|---:|
| ALT AUC + time | 7.67 | 5.6 to 10.0 |
| miR-122 AUC + time | 9.98 | 7.7 to 12.5 |
| ALT AUC + miR-122 AUC + time | 8.21 | 6.0 to 10.7 |

Additional serial-ALT features did not help either (Table S3). No added input lowered out-of-sample error below the single-biomarker baseline: collinear miR-122 and extra serial-ALT features added error.

**Table S3** Attempts to improve the model by adding inputs (tensor-product baseline, for internal comparability across this panel – not the additive GAM which achieved MAE 7.7%).

| Model inputs | LOOCV MAE (%) |
|---|---:|
| ALT AUC + time (baseline) | 7.9 |
| ALT AUC + miR-122 AUC + time | 8.4 |
| ALT AUC + time + time-of-peak-ALT | 7.9 |
| ALT AUC + miR-122 AUC + time-of-peak-ALT | 8.8 |

##### *4.1 The regeneration axis*

The ALT-AUC-to-necrosis relationship decays with time since dosing (Section 2.4), and regeneration is an obvious candidate explanation: as damaged tissue is repaired, a given cumulative enzyme exposure corresponds to less residual necrosis. We therefore asked whether representing regeneration improves prediction, either directly from proliferation or through a proxy obtainable without killing the animal. Neither did, which is why the model reported in the main text uses time since dosing alone. Proliferation is measured only at cull and so could never serve as a non-invasive input; it is included here as the biological check on that negative result.

Proliferation was assessed by 5-bromo-2'-deoxyuridine (BrdU) incorporation, administered approximately 1 h before a scheduled cull, on a multiplex immunofluorescence panel counterstained with DAPI and co-stained for HNF4α (hepatocytes) and ERG (endothelium). Whole-slide images were phenotyped by InForm trainable segmentation (Akoya Biosciences), classifying each nucleus by lineage and as BrdU-positive or negative. Because BrdU must be given approximately 1 h before cull, proliferation was available only for the 32 scheduled-cull dosed animals and the 5 controls and, like necrosis, only at cull. It is used as biological evidence on the regeneration axis, never as a model input.

Hepatocyte and endothelial BrdU incorporation was negligible until approximately 30 h after dosing and then rose (Fig. S6). A logistic form was supported for endothelial proliferation (inflection approximately 51 h, $R^2$ = 0.53), while hepatocyte proliferation showed a delayed rise with large between-animal heterogeneity at 36 to 48 h. Whole-tissue proliferation was uncorrelated with necrosis (r = minus 0.01, n = 32); resolving by lineage, hepatocyte proliferation correlated with time since dosing (r = 0.61) whereas endothelial proliferation did not, identifying hepatocyte regeneration as the time-dependent process.

Because a population regeneration trajectory is a deterministic function of time, and time is already a model input, it carries no independent information (regeneration index against time, r = 0.93). Entered as an extra covariate, or used to replace the model time axis, it made leave-one-animal-out prediction worse rather than better (8.8 to 10.1% against the 7.7% baseline): BrdU incorporation in this cohort was negligible before approximately 30 h, so a regeneration-time function carries little information across the 8 to 24 h window in which necrosis accrues, making it a degraded time axis. This may be a limitation of the method used - a one-hour BrdU pulse marks only nuclei in S phase during that hour, whereas markers of cell-cycle entry such as PCNA and Ki67 have been reported to rise considerably earlier after acetaminophen injury (Bhushan et al. 2014). Earlier cell-cycle entry would not change the result reported here, because the deployable predictor is derived from serial ALT rather than from proliferation, but it does mean the 30 h figure should be read as the limit of detection of this assay and not as the start of repair.

Per-animal proliferation does carry a late-time signal, but it is terminal. Within animals culled at 36 h or later, hepatocyte proliferation was negatively associated with residual necrosis (r = minus 0.64, against minus 0.26 before 36 h): among animals with similar cumulative ALT exposure, those further into regeneration had cleared more necrotic tissue. Entered as a terminal covariate it did not significantly improve prediction (paired reduction 0.8 percentage points, Wilcoxon p = 0.22).

The deployable, serial-ALT-derived proxy for this axis is the time of peak ALT. Adding it did not lower cross-validated MAE (7.67% to 8.19%, n = 49) and if anything slightly increased it (paired change minus 0.5 percentage points; 95% CI minus 1.2 to plus 0.1; Wilcoxon p = 0.09), and it could not replace time since dosing. It is reported as a lead for larger cohorts, not a confirmed gain.

A full black-box model, including informative missingness, did not help. Necrosis in the proliferation-absent (humane-endpoint) animals (mean 34.8%) did not differ from the proliferation-present animals (35.3%; Mann-Whitney p = 0.99). A histogram gradient-boosting regressor given cumulative ALT AUC, time, the four proliferation columns and, in a second variant, a missingness indicator, achieved 8.2% and 8.0% cross-validated error, not improving on cumulative ALT AUC and time alone (gradient boosting 8.0%, GAM 7.7%). Even a flexible learner given every terminal covariate and the missingness structure did not beat a single biomarker's kinetics.

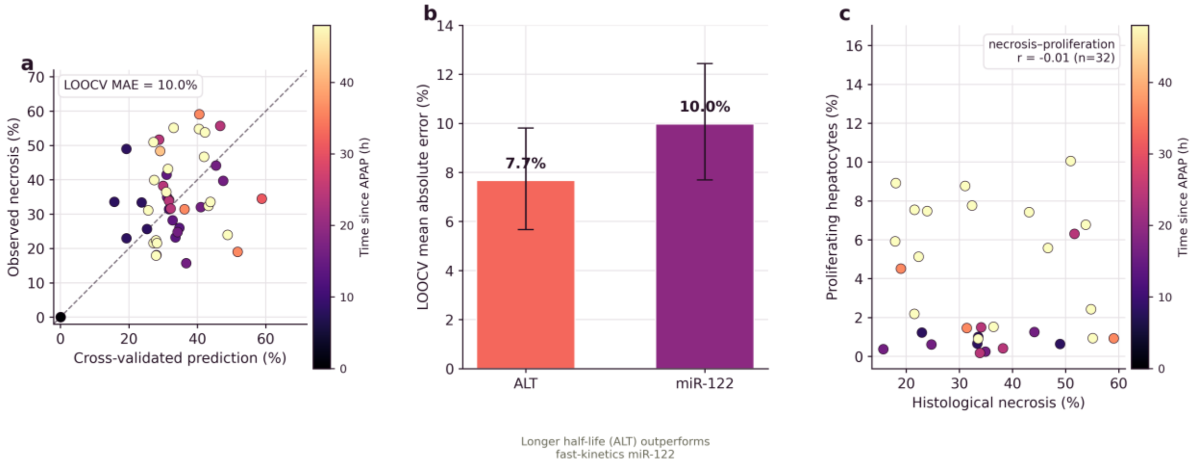


Fig. S5 Biomarker comparison. a miR-122 virtual histology by leave-one-animal-out cross-validation (MAE 10.0%). b ALT outperforms the faster-kinetics miR-122 by 2.3 percentage points. c Whole-tissue proliferation is uncorrelated with necrosis (r = minus 0.01); resolving by cell type, hepatocyte proliferation tracks time since dosing, indexing regeneration rather than the extent of necrosis. Error bars in b are 95% bootstrap confidence intervals.

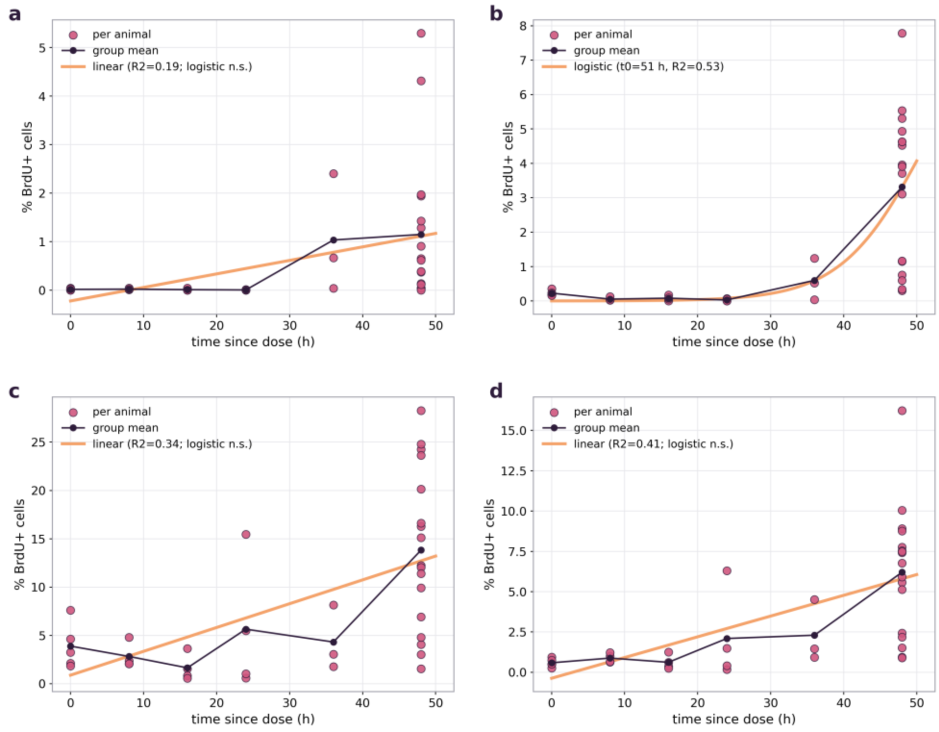

Fig. S6 Temporal morphology of proliferation, measured by BrdU incorporation and cross-sectional at cull. Each panel plots the percentage of BrdU-positive nuclei against time since dosing: a hepatocytes (HNF4α-positive), b endothelium (ERG-positive), c other or unclassified nuclei, and d all nuclei combined. Points are individual animals, the dark line joins group means and the orange line is the fitted trend, whose form and goodness of fit are given in each panel key. Hepatocyte and endothelial BrdU incorporation is negligible until approximately 30 h and then rises. Endothelial proliferation is the only lineage for which a logistic form is supported (inflection approximately 51 h, $R^2 = 0.53$); the hepatocyte, residual and total fractions rise gradually and are fitted linearly.

#### Section 5. Design, variability and the animal-number derivation

The main text states only the conclusion of this analysis. The derivation, the variance comparison between sampling designs and the worked example are given here in full, because they concern study design rather than the toxicological result.

##### *5.1 Sample-size derivation*

A two-group study aims to detect an absolute reduction $\delta$ in necrosis fraction with 80% power at $\alpha = 0.05$ (two-sided). Per-group sample size is $n = 2(z(1 \text{ minus } \alpha/2) + z(1 \text{ minus } \beta))^2\sigma^2/\delta^2$ (Festing and Altman 2002). The terminal design requires an independent cohort at each of T timepoints, giving a total of $2 \cdot T \cdot n(\text{terminal})$; the serial design measures all timepoints in one cohort, giving a total of $2 \cdot n(\text{serial})$. We used the measured pooled within-timepoint standard deviations: histological necrosis $\sigma = 0.120$ and model-predicted necrosis $\sigma = 0.108$ (leave-one-animal-out surrogate-error SD 0.115). Because these are close, the saving is dominated by the structural factor 1 minus 1/T and is robust to the precision assumption (Table S4).

**Table S4** Animals required by a terminal and a serial design across effect sizes and numbers of timepoints.

| Effect (Δ necrosis) | Timepoints | n/group terminal | n/group serial | Total terminal | Total serial | Reduction |
|---|---|---|---|---|---|---|
| 5% | 2 | 92 | 74 | 368 | 148 | 60% |
| 5% | 3 | 92 | 74 | 552 | 148 | 73% |
| 5% | 4 | 92 | 74 | 736 | 148 | 80% |
| 5% | 5 | 92 | 74 | 920 | 148 | 84% |
| 10% | 2 | 24 | 20 | 96 | 40 | 58% |
| 10% | 3 | 24 | 20 | 144 | 40 | 72% |
| 10% | 4 | 24 | 20 | 192 | 40 | 79% |
| 10% | 5 | 24 | 20 | 240 | 40 | 83% |
| 15% | 2 | 12 | 10 | 48 | 20 | 58% |
| 15% | 3 | 12 | 10 | 72 | 20 | 72% |
| 15% | 4 | 12 | 10 | 96 | 20 | 79% |
| 15% | 5 | 12 | 10 | 120 | 20 | 83% |
| 20% | 2 | 7 | 6 | 28 | 12 | 57% |
| 20% | 3 | 7 | 6 | 42 | 12 | 71% |
| 20% | 4 | 7 | 6 | 56 | 12 | 79% |
| 20% | 5 | 7 | 6 | 70 | 12 | 83% |

The worked example used in the main text is the 10% effect at three timepoints: 144 animals for the terminal design against 40 for the serial design. The main text quotes 40 to 48 because the range covers the precision assumption used for the predicted rather than the histological endpoint.

##### *5.2 Demonstrability of the variance-stability advantage*

Whether the serial design's variance advantage is demonstrable at this sample size was tested by bootstrap resampling (4,000 draws) of the ALT AUC coefficient of variation (CV) at each timepoint, for the serially-bled cohort ($n = 17$) against the independent cull groups ($n = 4$ to 6). The cull-group CV is estimated far less precisely, and propagating this into a sample-size calculation (to detect a 30% relative change at 80% power) shows the cull-based plan is unstable whereas the serial-bleed estimate is consistent (Table S5).

**Table S5** Bootstrap precision of the ALT AUC coefficient of variation by design.

| t (h) | Bleed CV % | Bleed 95% CI | Cull CV % | Cull 95% CI | n/grp (bleed) | n/grp (cull 95% CI) |
|---|---|---|---|---|---|---|
| 8 | 34 | 22 to 43 | 15 | 4 to 20 | 21 | 8 to 10 |
| 16 | 37 | 24 to 46 | 26 | 10 to 32 | 26 | 3 to 20 |
| 24 | 38 | 26 to 46 | 31 | 5 to 43 | 27 | 10 to 33 |
| 36 | 37 | 27 to 44 | 45 | 8 to 55 | 26 | 3 to 55 |

Variance-ratio F-tests between designs are non-significant at this n ($p = 0.30$ to 1.00). The demonstrable effect is therefore the precision of the variance estimate, which is driven by group size, and not a proven difference in the underlying variances. A study powered from cull groups could by chance call for as few as 3 or as many as 55 animals per arm, where the serial cohort gives a stable 21 to 27.

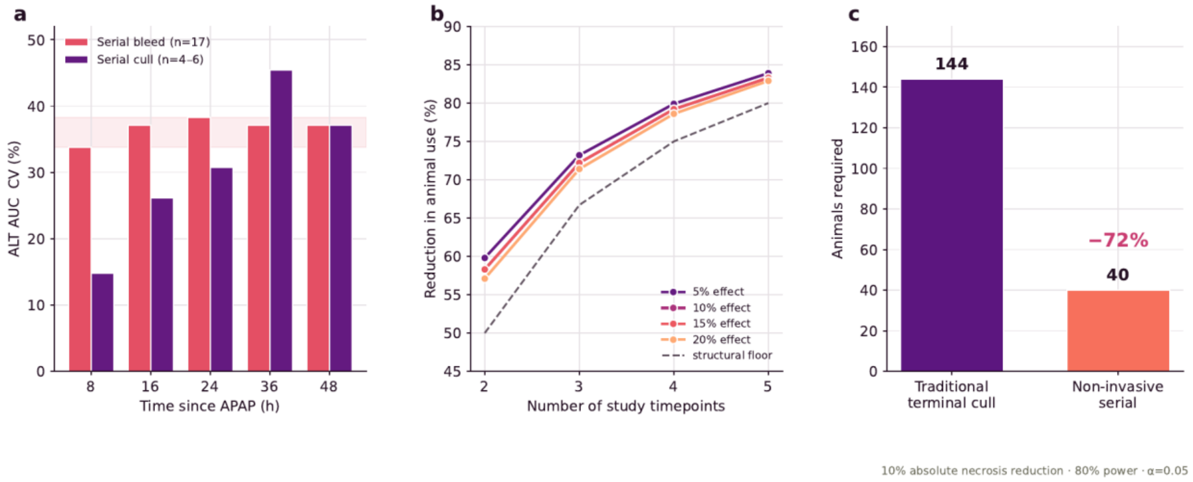


Fig. S7 Design and animal-number consequences. a Sampling-design variability: coefficient of variation of cumulative ALT AUC by timepoint, stable for serial bleeding (34 to 38%; the shaded band marks this range) and variable for the independent cull groups (15 to 45%); the point CVs do not differ significantly (variance-ratio F-tests $p = 0.30$ to 1.00). b Projected animal saving: the reduction in animal use achieved by a serial design relative to a terminal design (higher is better) against the number of study timepoints, for four target effect sizes. The dashed line is the structural floor, 100(1 minus 1/T): a serial design uses one cohort in place of one per timepoint, so at equal per-group precision it saves 1 minus 1/T of the animals. The effect-size lines sit just above this floor because the model-predicted endpoint had marginally lower within-timepoint variance than histology (SD 0.108 versus 0.120), and the spread across effect sizes is a sample-size-rounding effect. c Worked example: a two-group, three-timepoint study powered on a 10-percentage-point necrosis reduction needs 144 animals on a terminal design against 40 on a serial design using the predicted-endpoint variance, a 72% reduction. The main text quotes 40 to 48 because the upper bound uses the histological rather than the predicted variance.

#### Section 6. Per-animal data

Data for the 44 dosed mice that entered the models. Cumulative ALT AUC is given in $\times10^3$ $U \cdot L^{-1} \cdot h$; necrosis is percentage area from the scoring output identified in Section 1.3; the prediction is the leave-one-animal-out estimate from the additive GAM. The five fasted controls had 0% necrosis at $t = 0$ and are not tabulated. The full analysis dataset, including miR-122 and proliferation columns, accompanies this ESM as a spreadsheet.

| Mouse | Cull (h) | ALT AUC | Necrosis % | Predicted % |
|---|---|---|---|---|
| 206 | 8 | 42.2 | 49 | 28.2 |
| 207 | 8 | 37.5 | 33.4 | 25.1 |
| 209 | 8 | 40.6 | 33.6 | 27.2 |
| 210 | 8 | 33 | 23 | 22.1 |
| 211 | 16 | 111.8 | 24.8 | 37.2 |
| 212 | 16 | 145.8 | 44.1 | 48.6 |
| 213 | 16 | 87.8 | 15.7 | 29.2 |
| 214 | 16 | 76.6 | 34.9 | 25.5 |

| Mouse | Cull (h) | ALT AUC | Necrosis % | Predicted % |
|---|---|---|---|---|
| 215 | 14 | 104.5 | 41.5 | 39.8 |
| 216 | 24 | 87.8 | 34.1 | 19.5 |
| 217 | 24 | 144.3 | 33.9 | 32 |
| 218 | 24 | 151.8 | 51.7 | 33.6 |
| 219 | 24 | 134.6 | 31.6 | 29.8 |
| 220 | 24 | 134.7 | 38.2 | 29.8 |
| 221 | 16 | 142.2 | 39.7 | 47.4 |
| 222 | 8 | 49.1 | 25.6 | 32.9 |
| 223 | 36 | 299.7 | 59.1 | 44.1 |
| 224 | 36 | 132.5 | 31.4 | 19.5 |
| 225 | 36 | 161.6 | 19 | 23.8 |
| 226 | 11 | 89.7 | 31.4 | 43.6 |
| 227 | 48 | 369.6 | 32.4 | 40.8 |
| 228 | 48 | 418.4 | 55.1 | 46.1 |
| 229 | 42 | 401.9 | 48.4 | 50.7 |
| 230 | 48 | 479.9 | 33.5 | 52.9 |
| 231 | 48 | 502.5 | 36.4 | 55.4 |
| 232 | 13 | 94.8 | 23.2 | 38.9 |
| 233 | 24 | 226.5 | 55.7 | 50.2 |
| 234 | 13 | 101 | 26 | 41.5 |
| 235 | 48 | 387.9 | 53.8 | 42.8 |
| 236 | 48 | 274.8 | 54.8 | 30.3 |
| 237 | 48 | 200 | 18 | 22 |
| 238 | 14 | 109 | 28.2 | 41.5 |
| 239 | 48 | 176.1 | 21.5 | 19.4 |
| 240 | 31 | 308.7 | 34.5 | 52.9 |
| 241 | 48 | 271.7 | 43.1 | 30 |
| 242 | 16 | 145 | 32 | 48.3 |
| 243 | 48 | 273.7 | 39.9 | 30.2 |
| 244 | 48 | 380.8 | 46.7 | 42 |
| 245 | 48 | 221.2 | 31.1 | 24.4 |
| 246 | 48 | 182.3 | 21.6 | 20.1 |
| 247 | 48 | 247.6 | 23.9 | 27.3 |
| 248 | 48 | 140.6 | 17.9 | 15.5 |
| 249 | 48 | 197 | 22.3 | 21.7 |
| 250 | 48 | 303.8 | 51 | 33.5 |
