## Supplementary material for "Non-invasive quantification of hepatic necrosis from circulating alanine aminotransferase kinetics in acetaminophen-treated mice in vivo": ARRIVE

### ARRIVE 2.0 reporting checklist

#### The ARRIVE Essential 10

| # | Item | What is reported | Where |
|---|---|---|---|
| 1 | Study design | Single in vivo experiment. One experimental group (350 mg/kg APAP i.p.) sampled serially and culled at seven scheduled timepoints, plus an undosed fasted control group at t = 0. The experimental unit is the individual animal. NOTE: the manuscript no longer carries an explicit statement that the results were not reproduced in an independent experiment; see the note below the tables. | p. 2, l. 48–53; Suppl. S1.1 |
| 2 | Sample size | 50 mice acquired, 45 dosed and 5 undosed controls; 1 excluded, giving n = 49 analysed. Group sizes 5 to 17 per timepoint, given in full in the Supplementary Information. No a priori power calculation: this is a proof-of-concept calibration study, and the sample-size implications derived from it are reported in Suppl. Section 5. | p. 2, l. 53; Suppl. S1.1, S5 |
| 3 | Inclusion and exclusion criteria | Pre-specified: animals not surviving to a scheduled or humane-endpoint cull with paired biomarker and histology data were excluded. One mouse found dead at 7 h was excluded. Seven animals reaching a humane endpoint off schedule were retained and analysed at the nearest scheduled timepoint. No other exclusions; no data points were removed from any analysis. | p. 2, l. 52–53; Suppl. S1.1 |
| 4 | Randomisation | Animals were NOT randomised to cull timepoint. Allocation was sequential because animals were co-housed by timepoint to maintain social housing and reduce isolation stress. This is a limitation and is stated as such. | Suppl. S1.1 |
| 5 | Blinding | Plasma ALT and histological necrosis were measured blind to group. Allocation itself was not concealed, since cull timepoint is inherent to the procedure. | p. 3, l. 55–56; Suppl. S1.1 |
| 6 | Outcome measures | Primary outcome: percentage centrilobular necrotic area on haematoxylin and eosin whole-slide images, quantified by trainable tissue segmentation. Predictor: cumulative plasma ALT AUC to cull and time since dosing. Performance metric: leave-one-animal-out cross-validated mean absolute error. | p. 3, l. 56–58; Suppl. S1.3 |
| 7 | Statistical methods | Generalised additive model with one penalised smooth per predictor on the square-root response scale; leave-one-animal-out cross-validation with smoothing parameters selected inside each training fold; 10,000-sample bootstrap 95% confidence intervals. Comparator panel evaluated identically. The R and Python cross-implementation is reported in Suppl. S2.2. | p. 3, l. 59–61; Suppl. S2, S3 |
| 8 | Experimental animals | Mus musculus, C57BL/6J, male, twelve weeks old, Charles River UK. Fasted 14 h before dosing. | p. 2, l. 48; Suppl. S1.1 |
| 9 | Experimental procedures | Single intraperitoneal dose of 350 mg/kg acetaminophen in warm sterile saline. Dose justified as producing reproducible submaximal centrilobular necrosis within a UK moderate-severity limit, against the 500 mg/kg more commonly used elsewhere. Serial 40 µL tail-vein sampling at baseline and 8, 16, 24, 36 and 48 h until cull, with a terminal cardiac sample. Liver fixed, sectioned at 4 µm, H&E stained. | p. 2, l. 48–51; p. 3, l. 55–58; Suppl. S1.2, S1.3 |
| 10 | Results | Necrosis 0 to 59%, cohort mean 32%. Cross-validated MAE 7.7% necrotic area (95% CI 5.6 to 10.0), n = 49. Full comparator panel with confidence intervals and per-animal data in the Supplementary Information. | p. 3, l. 67–74; Suppl. Table S1, Section 6 |

#### The Recommended Set

| # | Item | What is reported | Where |
|---|---|---|---|
| 11 | Abstract | Structured as a single paragraph stating background, species, dose, methods, primary result with confidence interval and conclusion. | p. 2, l. 19–32 |
| 12 | Background | Relevance to toxicology stated: histology is terminal, biomarkers are uncalibrated, and the kinetic information needed is already collected and discarded. | p. 2, l. 36–40 |
| 13 | Objectives | Whether cumulative ALT exposure and time since dosing are together sufficient to estimate histological necrosis. | p. 2, l. 41–44 |
| 14 | Ethical statement | Approved by the University of Edinburgh Animal Welfare and Ethical Review Body, project licence 093-LFR-24, under the Animals (Scientific Procedures) Act 1986. | p. 2, l. 46–47; p. 4, l. 113–115 |

| # | Item | What is reported | Where |
|---|---|---|---|
| 15 | Housing and husbandry | Group-housed five per cage, 12 h light/dark cycle, free access to water, 14 h fast before dosing. | Suppl. S1.1 |
| 16 | Animal care and monitoring | Welfare assessed against protocol-specific scoring criteria; seven animals reached a humane endpoint and were culled off schedule. One animal was found dead at 7 h. | p. 2, l. 52; Suppl. S1.1 |
| 17 | Interpretation and scientific implications | Result is biologically coherent; limitations of mechanism, dose, sex and strain are stated, as is the assumption that injury releases ALT proportionally to hepatocyte death. | p. 3, l. 89 to p. 4, l. 95 |
| 18 | Generalisability and translation | Stated as untested beyond one mechanism, dose, sex and strain; external validation named as the necessary next step. Principle is not liver-specific. | p. 3, l. 90 to p. 4, l. 95 |
| 19 | Protocol registration | The study protocol was not pre-registered. No analysis plan was registered. | Not applicable |
| 20 | Data access | Per-animal data for all 44 dosed animals are provided in the Supplementary Information and as a spreadsheet. Analysis code available from the corresponding author. | p. 4, l. 121–122; Suppl. Section 6 |
| 21 | Declaration of interests | Declared under Statements and Declarations. Funding sources listed with a statement that funders had no role in design, analysis or the decision to publish. | p. 4, l. 105–112 |
